# Comparison of cell-cycle gene expression dynamics and mRNA kinetics across mouse and human pluripotent systems

**DOI:** 10.64898/2026.09.14.751360

**Authors:** Maulik K. Nariya, David Santiago-Algarra, Gianni Zanardelli, Idris Kouadri Boudjelthia, Tao Ye, Christelle Thibault-Carpentier, Sophie Jarriault, Nacho Molina

**Affiliations:** Université de Strasbourg, CNRS, INSERM, IGBMC UMR 7104-UMR-S 1258, F-67400 Illkirch, France; Theoretical and Scientific Data Science, Scuola Internazionale Superiore di Studi Avanzati (SISSA), Trieste, Italy; School of Science and Technology, IE University, Madrid, Spain

## Abstract

Cell-cycle remodeling is fundamental to pluripotency and lineage commitment, yet whether its transcriptional and post-transcriptional architecture is conserved across species and developmental states has remained unresolved. Here we introduce Ciclopes, a biology-informed deep-learning framework that resolves continuous cell-cycle phase and phase-dependent mRNA transcription and degradation directly from single-cell RNA sequencing. Applying Ciclopes across six mouse and human pluripotent stem-cell systems spanning naїve and primed states, we uncover striking divergence in transcriptional complexity and oscillatory control: mouse systems sustain elevated baseline expression of core cell-cycle regulators, while human systems trade higher baseline expression for larger oscillatory amplitude. Strikingly, mRNA degradation timing remain far more conserved across systems than transcription timing, exposing post-transcriptional regulation as a stable evolutionary backbone. As human iPSCs differentiate into definitive endoderm, cells progressively exit the cell cycle, cell-cycle-coupled gene networks contract, and surviving regulators oscillate with larger amplitude. Ciclopes establishes a general framework for dissecting how pluripotent cells tune proliferation across evolutionary and developmental transitions.

## Introduction

Pluripotent stem cells combine rapid self-renewal with developmental competence, and transitions between pluripotent states are accompanied by marked changes in cell-cycle organization. Many naїve and rapidly pro-liferating pluripotent states exhibit an abbreviated G1 phase and a high proportion of cells in S phase, whereas progression toward primed pluripotency and lineage commitment is accompanied by reinforcement of G1 control and extensive remodeling of proliferative behavior [1–4]. Developmental competence also changes as cells progress through the cell cycle: human pluripotent cells in early and late G1 differ in their signaling responses and propensity to adopt alternative lineages [5]. Thus, rather than operating as an autonomous program of cell proliferation, cell-cycle progression is tightly integrated with the signaling and gene-regulatory networks that govern pluripotency and lineage commitment. Yet pluripotent states differ substantially across species and developmental contexts in their molecular identity, culture conditions, and proliferative properties [6–9]. It therefore remains unclear which features of cell-cycle regulation are shared across pluripotent systems and which are reorganized in a context-dependent manner.

Resolving this question requires going beyond discrete phase assignments to characterize the continuous molecular programs that accompany progression through the cell cycle. DNA-content and nucleotide-incorporation assays, together with live-cell reporters, have established major differences in phase structure and cell-cycle progression across pluripotent states, but do not by themselves resolve genome-wide transcriptional programs. Conversely, single-cell RNA sequencing captures genome-wide expression in large populations of asynchronously cycling cells, from which continuous cell-cycle progression can be reconstructed [10]. Importantly, phase-dependent mRNA abundance reflects the combined action of transcription and post-transcriptional processes, including RNA processing and degradation. Consequently, similar expression dynamics can arise from different underlying kinetic programs, while changes in transcript abundance do not directly reveal whether transcriptional or post-transcriptional regulation has been remodeled. Recent phase-resolved modeling in mouse embryonic stem cells showed that transcription, RNA processing, and degradation are themselves dynamically regulated across the cell cycle [11]. Whether the temporal organization of these processes is conserved across pluripotent systems, however, remains unknown.

Several computational approaches have been developed to infer continuous cell-cycle progression from single-cell transcriptomes. Cyclum uses an autoencoder to recover latent periodic structure, whereas DeepCycle exploits spliced and unspliced transcript abundances to infer continuous phase and oscillatory gene-expression dynamics. VeloCycle formulates cell-cycle progression within a probabilistic, manifold-constrained RNA-velocity framework, while FourierCycle extends phase-resolved modeling to multiple components of mRNA metabolism [10–14]. These approaches differ in how they incorporate prior biological information and in whether phase inference depends on spliced and unspliced measurements. For comparative analyses across heterogeneous biological systems, a useful framework should infer a context-specific, biologically oriented phase directly from gene expression, while allowing phase-resolved kinetic modelling to be incorporated when spliced and unspliced measurements are available.

Here, we introduce Ciclopes, a biology-informed autoencoder that jointly infers continuous cell-cycle phase and phase-dependent gene-expression dynamics. Ciclopes first positions cells in an S–G2/M manifold defined by established S-phase and G2/M-associated transcriptional programs, providing a biologically oriented initial ordering of the cell cycle. A neural-network encoder subsequently maps each transcriptome onto a one-dimensional circular phase, while an interpretable Fourier decoder reconstructs periodic gene-expression profiles and jointly refines the inferred ordering. Phase inference depends only on gene-expression measurements; when spliced and unspliced transcript abundances are available, the inferred phase can additionally be coupled to a kinetic model to estimate effective phase-dependent transcription and degradation profiles. We applied Ciclopes to six single-cell RNA-sequencing datasets representing mouse and human pluripotent systems spanning naїve and primed states, and to a time course of human induced pluripotent stem-cell differentiation toward definitive endoderm. We used these data to ask how cell-cycle composition and oscillatory gene-expression programs vary across pluripotent contexts, whether transcriptional and post-transcriptional dynamics exhibit comparable cross-system organization, and how these programs are remodeled during lineage commitment.

Across the analyzed systems, we observed distinct quantitative regimes of cell-cycle-associated expression. Mouse datasets generally showed higher relative baseline expression of core cell-cycle regulators, whereas several human datasets displayed larger phase-dependent oscillatory amplitudes. Despite greater variation in inferred transcriptional timing, degradation profiles were substantially more concordant across systems, suggesting that the temporal organization of mRNA degradation represents a comparatively stable component of pluripotent cell-cycle regulation. During differentiation toward definitive endoderm, the proportion of cells with low cycling activity increased, the actively cycling population became enriched in G1, and fewer genes remained coupled to cell-cycle progression, while several remaining regulators exhibited increased phase-dependent amplitude. Together, these results support a model in which context-dependent transcriptional programs are superimposed on a more stable post-transcriptional architecture, and provide a framework for resolving continuous cell-cycle regulation across pluripotent states and developmental transitions.

## Results

### Summary statistics reveal biological differences in the transcriptional complexity of pluripotent systems

To compare cell cycle dynamics across pluripotent systems, we examined single-cell RNA-seq (scRNA-seq) profiles from six pluripotent systems treated with different culture conditions: mouse embryonic stem cells (mESCs), mouse epiblast stem cells (mEpiSCs), human naїve embryonic stem cells (hNES1), human embryonic stem cells (hESCs), and human induced pluripotent stem cells (hiPSCs) [10, 15–18]. mESCs and hNES1 are considered to resemble the naїve pluripotent state, corresponding to the pre-implantation epiblast, whereas mEpiSCs, hESCs, and hiPSCs resemble the primed pluripotent state, corresponding to the post-implantation epiblast [6, 7, 9, 19]. Fig. 1a shows the distribution of raw counts per cell (see **Methods** for data processing details); counts were aproximately log-normally distributed in all systems, the colored numbers indicate the median counts observed in each condition. Fig. 1b plots the coefficient of variation, i.e. the relative noise, against mean expression for all genes across conditions. As expected, genes with lower mean expression showed higher noise, while genes with higher mean expression showed lower noise, with noise saturating to roughly 0.1–0.9 at a mean expression of approximately 100 counts per cell in most conditions. Fig. 1c shows the number of genes detected as a function of total counts per cell; unsurprisingly, genes detected increased with total counts, saturating around 10,000–12,000 genes at 75,000–100,000 total counts per cell. For clarity, panels 1b and 1c are also shown broken out by individual model system in Fig. S1.

**Figure 1.**
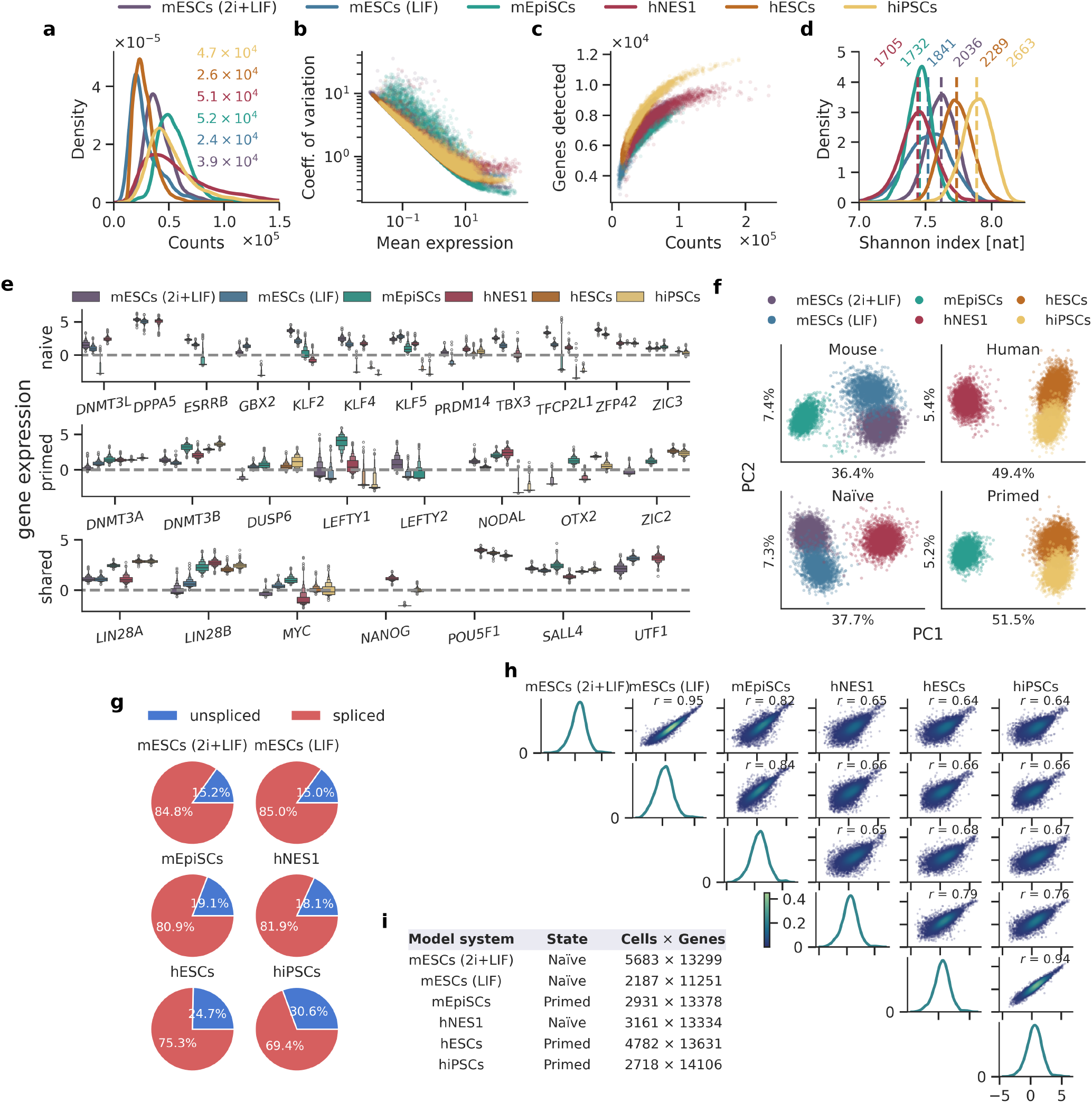
Summary statistics and expression levels of pluripotency markers across model systems. **a** Distribution of raw mRNA counts per cell across the six pluripotent systems. Colored numbers indicate the median counts per condition. **b** Coefficient of variation (*σ/µ*) versus mean expression for all genes across conditions, illustrating the characteristic decline in relative noise with increasing mean expression. **c**. Number of genes detected versus total counts per cell across conditions. Individual panels for each system are shown in Fig. S1. **d** Distribution of Shannon’s diversity index across cells in each system; dashed vertical lines and colored numbers indicate the mean diversity index per condition. **e** Sanity-normalized, mean-centered, expression of naїve (top), primed (middle), and shared (bottom) pluripotency marker genes across the six systems. The dashed grey line at 0 indicates the mean expression across all genes and all cells. Mouse genes were mapped to human homologs using the Mouse Genome Informatics database. **f** PCA of combinations of datasets–top left: mESCs (2i+LIF), mESCs (LIF), mEpiSCs, top right: hNES1, hESCs, hiPSCs, bottom left: mESCs (2i+LIF), mESCs (LIF), hNES1, and bottom right: mEpiSCs, hESCs, hiPSCs. **g** Percentage of spliced and unspliced mRNA per cell in each system. **h** Pairwise scatterplots (lower triangle) and density distributions (diagonal) of pseudobulked mean gene expression across the six systems; Pearson correlation coefficients (*r*) are indicated for each pairwise comparison. **i** Cells *×* genes dimensions and developmental state (naїve/primed) for each of the six model systems used in this study.

Finally, we used Shannon’s diversity index or Shannon index (see **Methods**) to quantify the information content of individual cells. Note, to account for the differences in the depth, we calculated the Shannon index using 8166 overlapping genes between the six systems. Shan-non indices ranged from approximately 7–8.25 nats across the pluripotent systems. hNES1 had the lowest mean index of all systems, followed by mEpiSCs and mESCs (LIF)—likely reflecting the differences noise structure of these data. By contrast, mESCs (2i+LIF), hESCs, and hiPSCs had comparable noise structure, yet showed distinct Shannon indices, suggesting genuine biological differences in transcriptional complexity arising from species differences as well as from naїve versus primed identity.

We compared the gene expression levels of naїve and primed pluripotency markers [20, 21]. To enable comparison across models, we mean-centered the Sanity-corrected values (the distribution of Sanity-corrected, mean-centered expression across cells is shown in Fig. S2). Fig. 1e shows mean-centered expression of pluripotency markers across the model systems; the dashed grey line at 0 represents the mean expression across all genes and all cells in the dataset. To enable cross-species comparison, mouse genes were mapped to their human homologs using the Mouse Genome Informatics database [22]. As expected, naїve markers were expressed at higher levels in the naїve-state systems, mESCs and hNES1, while primed markers were expressed at higher levels in the primed-state systems, mEpiSCs, hESCs, and hiPSCs; shared markers were expressed at comparable levels across all systems. However, several markers showed unexpected patterns: *KLF2* levels in hNES1 were similar to those in mEpiSCs, *PRDM14* levels in mESCs were similar to those in hESCs and hiPSCs, and *ZFP42* was highly expressed across all pluripotent systems, albeit at higher levels in mESCs. The interquartile ranges were narrow for most pluripotency markers, indicating low cell-to-cell variability in their expression. *LEFTY1* and *LEFTY2* showed the largest variation in expression across systems; notably, *LEFTY1* levels were significantly higher in mEpiSCs than in hESCs or hiPSCs, despite all three representing the primed state.

Fig. 1f shows principal component analysis performed in the space of 35 pluripotency markers [20, 21, 23– 28], applied separately to four combinations of systems: mouse systems alone (mESCs (2i+LIF), mESCs (LIF), mEpiSCs), human systems alone (hNES1, hESCs, hiP-SCs), naїve systems alone (mESCs (2i+LIF), mESCs (LIF), hNES1), and primed systems alone (mEpiSCs, hESCs, hiPSCs). In every combination, PC1 explained substantially more variance than PC2 (36–52% versus 5–7%) and consistently separated cell type or culture condition, whereas PC2 captured finer-grained, within-group heterogeneity. Rather than resolving into sharply separated clusters, cells occupied continuous, overlapping distributions along PC1, with no system forming an isolated island in marker space. This continuity echoes the marker-level exceptions noted above—*KLF2* expression in hNES1 tracking with mEpiSCs, *PRDM14* in mESCs tracking with hESCs and hiPSCs, and *ZFP42* remaining broadly expressed across all six systems—and together these observations argue that naїve and primed identity is better conceptualized as a continuum of pluripotent states than as two discrete, categorically separable programs, with the naїve/primed marker framework capturing gradients of identity rather than fixed boundaries [29].

Fig. 1g shows the percentage of unspliced and spliced mRNA—relevant for RNA velocity modeling—across cells in each pluripotent system. Human systems tended to show higher fractions of unspliced mRNA than mouse systems. Within each species, primed-state systems (mEpiSCs in mouse; hESCs and hiPSCs in human) showed slightly elevated unspliced mRNA fractions relative to their naїve counterparts (mESCs and hNES1, respectively). Finally, Fig. 1h shows a pairwise comparison of pseudobulked mean gene expression across the pluripotent model systems. As expected, gene expression profiles in mESCs (2i+LIF) correlated strongly with those in mESCs (LIF) (*r* = 0.95), and hESC profiles correlated strongly with hiPSC profiles (*r* = 0.94). mEpiSCs also correlated well with mESCs (2i+LIF) and mESCs (LIF) (*r* = 0.82 and *r* = 0.84, respectively), though less strongly than the mESC-mESC comparison, consistent with naїve-versus-primed divergence. Correlations between the human systems (hNES1, hESCs, hiPSCs) and the mouse systems (mESCs, mEpiSCs) ranged from *r* = 0.64 to 0.66, while the correlation between hNES1 and hESCs was *r* = 0.79, and between hNES1 and hiPSCs was *r* = 0.76.

Taken together, these summary statistics establish that although the six pluripotent systems examined here share the core hallmarks of pluripotency, they are far from transcriptionally interchangeable and that species identity, culture condition, and developmental state each leave distinct and quantifiable imprints on transcriptional complexity, marker expression, and splicing dynamics. This heterogeneity argues against treating “pluripotent stem cells” as a monolithic reference state, and instead motivates a system-by-system dissection of the cell cycle dynamics that underlie these differences—the focus of the remainder of this study.

### Cell cycle regulators have higher mean expression in mouse and larger amplitude of oscillations in human pluripotent systems

We developed Ciclopes, a biology-informed deep learning tool for inferring continuous cell cycle phase and mRNA kinetics from scRNA-seq profiles (Fig. 2a). Ciclopes is an autoencoder in which a neural network encoder maps each cell’s gene expression to a one-dimensional circular latent variable, the inferred cell cycle phase *θ*, while a Fourier-based decoder reconstructs gene expression as a function of *θ*, capturing its oscillatory dynamics across the cycle. Phase inference is initialized using the S–G2/M manifold, a biology-informed estimate of cell cycle position built from the S-G2/M-phase marker scores (Fig. 2b), and *θ* is then iteratively refined jointly with the Fourier coefficients during training, anchored to this initial estimate to preserve biological grounding while allowing the phases to redistribute smoothly around the cycle Fig. S3 shows comparison of cell cycle phases inferred by Ciclopes, DeepCycle, VeloCycle.

**Figure 2.**
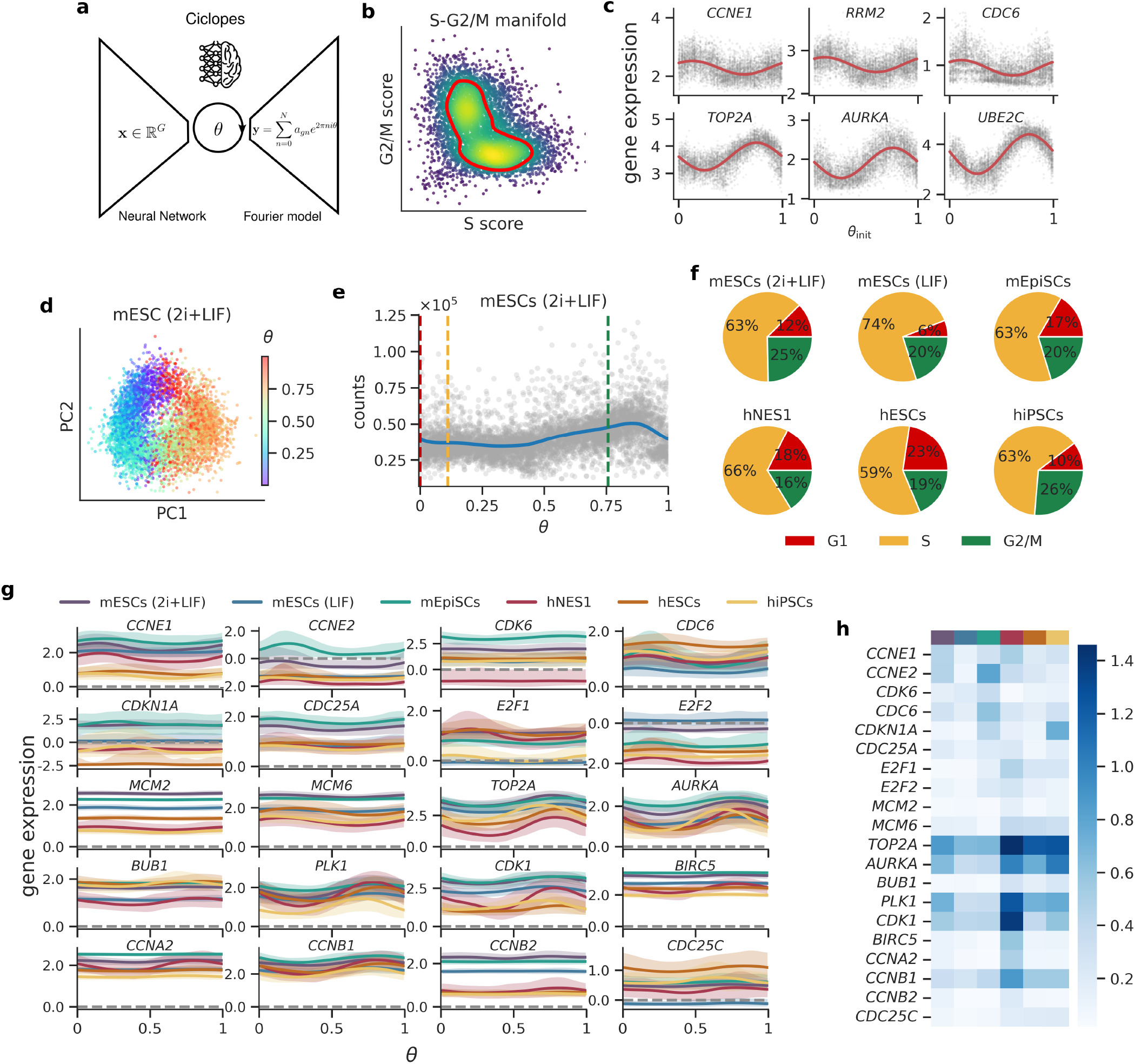
Cell cycle dynamics across pluripotent systems. **a** Schematic of the Ciclopes model architecture, an autoencoder in which the encoder is a neural network that maps gene expression x to a scalar cell cycle phase *θ*, and the decoder is a Fourier-based generative model that reconstructs gene expression as a function of *θ*. **b** Representative S–G2/M manifold used to obtain an initial estimate of cell cycle phase; color indicates local point density, and the red contour marks the estimated initial cell cycle trajectory. **c** Expression of representative cell cycle marker genes (*CCNE1, RRM2, CDC6, TOP2A, AURKA, UBE2C*) as a function of the initial phase estimate, *θ*_init_. Grey points are individual cells; red lines show the fitted Fourier-based expression model. **d** Cell cycle phase *θ* projected onto the top two principal components (PCs) of gene expression for mESCs (2i+LIF); color indicates *θ*. **e** Total mRNA counts per cell as a function of inferred cell cycle phase *θ* for mESCs (2i+LIF); dashed lines mark the inferred G1/S (yellow) and S/G2 (green) phase boundaries. Corresponding panels for all six systems are shown in Fig. S4. **f** Percentage of cells assigned to G1 (red), S (yellow), and G2/M (green) phases in each pluripotent system, based on transcriptional phase boundaries. **g** Expression dynamics of 20 representative cell cycle regulators as a function of *θ* across all six pluripotent systems; the solid lines indicate the moving average of the gene expression values where as the shaded regions indicate *±*2*σ* spread around the average gene expression. **h** Heatmap of oscillation amplitude for each cell cycle regulator (rows) across the six pluripotent systems (columns).

Fig. 2c shows expression of representative cell cycle markers (*CCNE1, RRM2, CDC6, TOP2A, AURKA, UBE2C*) as a function of the initial phase estimate, *θ*_init_, together with the fitted Fourier model. Fig. 2d projects the inferred phase *θ* onto the first two principal components of gene expression for mESCs (2i+LIF), showing that *θ* traces a continuous, biologically ordered trajectory through transcriptomic space. Fig. 2e shows total UMI counts per cell as a function of *θ* for the same system, with dashed lines marking the inferred G1/S and S/G2 boundaries (see **Methods** for phase boundary determination); corresponding panels for all six systems are shown in Fig. S4.

Fig. 2f shows the proportion of cells assigned to G1, S, and G2/M phases in each system. Human systems generally showed a higher proportion of cells in G1 than mouse systems. Within each species, the primed-state system had a higher G1 fraction than its naїve counterpart: mEpiSCs exceeded mESCs (2i+LIF and LIF), and hESCs exceeded hNES1. These findings are consistent with results in Boward et al, where the fractions of cells in different phases are determined by Edu-DNA content assay [1]. hiPSCs, however, did not follow this trend, showing the lowest G1 fraction of any human system despite representing the primed state. Note, the phase proportions here are derived from cell cycle boundaries defined using the transcriptional states of S and G2/M markers. Table S2 provides the cell cycle durations measured/inferred in the literature [2, 3, 30–37].

Fig. 2g shows expression dynamics of representative cell cycle markers across all six systems. Relative to background expression, several regulators—including *CCNE1, CDKN1A, CDC25A, MCM2, MCM6, TOP2A*, and *AU-RKA*—showed higher mean expression in mouse systems than in human systems. For a subset of genes, including *E2F1, PLK1, CCNB1*, and *CCNA2*, expression levels in hNES1 more closely resembled those of the mouse systems than the other human systems. As expected, most of these markers showed oscillatory expression across the cell cycle, but the amplitude of these oscillations varied considerably across systems. Fig. 2h summarizes this variation as a heatmap of oscillation amplitude for each cell cycle regulator across all six systems. Overall, mouse systems tended toward higher mean expression of cell cycle regulators, while human systems—particularly hNES1—tended toward larger amplitude oscillations in their expression dynamics.

Together, these results show that Ciclopes recovers a continuous, biologically consistent cell cycle trajectory across all six pluripotent systems, revealing that cell cycle regulators are tuned differently across species: mouse systems favor elevated baseline expression of core regulators, whereas human systems favor larger-amplitude oscillations around a lower baseline. This divergence in expression strategy, layered on top of species- and state-dependent differences in phase distribution, indicates that mouse and human pluripotent cells achieve cell cycle control through quantitatively distinct regulatory regimes rather than a single conserved expression program.

### Kinetics of post-transcriptional regulation remain conserved across the pluripotent systems

While the phase model recovers the oscillatory dynamics of gene expression as a function of cell cycle phase, it is agnostic to the underlying molecular processes that generate those dynamics. In particular, it cannot resolve the cell cycle dependence of transcription and degradation—the two processes that jointly determine mRNA abundance at each phase. To address this, we constructed a kinetic model that explicitly accounts for the steps involved in mRNA metabolism: transcription, splicing, and degradation.

The top two rows of Fig. 3a show heatmaps of genome-wide oscillatory gene expression dynamics, split into spliced and unspliced transcripts. Genes were filtered based on explained variance and fold-change in oscillation amplitude relative to baseline (see **Methods**); the number in parentheses above each panel indicates the number of genes surviving this criterion. In both species, the naїve systems—mESCs (LIF) and hNES1—tended to have more genes with oscillatory expression than their primed counterparts—mEpiSCs, hESCs, and hiP-SCs—with the exception of mESCs (2i+LIF), suggesting that a broader set of genes is coupled to cell cycle regulation earlier in development. We also observed distinct patterns in the “waves” of gene expression across the cell cycle, evident in both spliced and unspliced transcripts, pointing to biological differences in the allocation of transcriptional resources both between species and between naїve and primed states. Fig. S5 shows the corresponding enrichment analysis for genes reaching peak expression in G1 (red), S (yellow), and G2/M (green).

**Figure 3.**
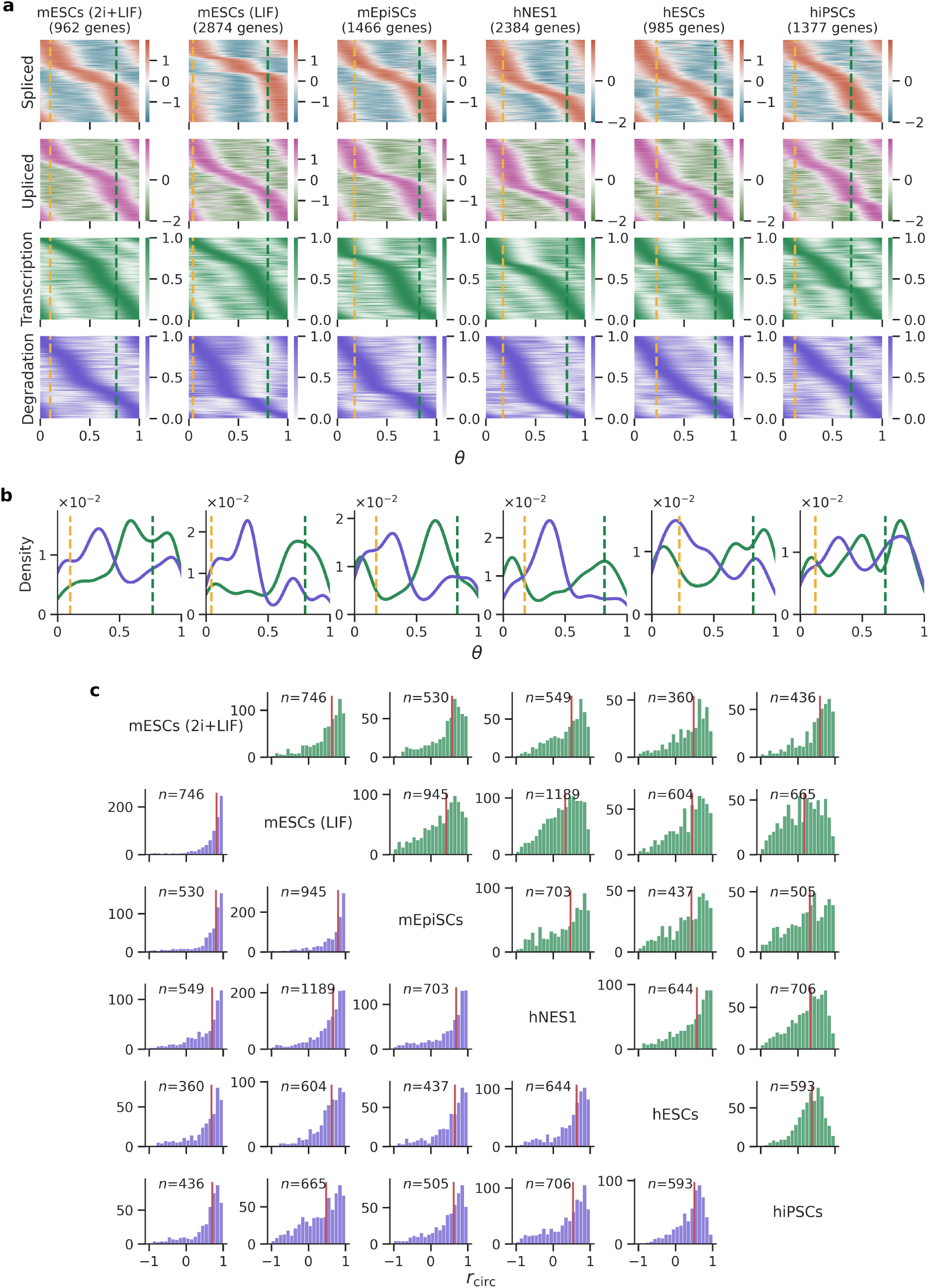
Kinetics of mRNA transcription and degradation during the cell cyle across pluripotent systems. **a** Heatmaps of cell cycle phase-ordered gene expression dynamics for genes passing explained-variance and amplitude filtering criteria (see **Methods**) in each of the six pluripotent systems (number of genes indicated above each column). From top to bottom: spliced expression, unspliced expression, inferred transcription rate, and inferred degradation rate, each scaled and ordered by the phase at which the gene reaches its peak value. Dashed yellow and green lines mark the inferred G1/S and S/G2 phase boundaries, respectively. **b** Density histograms of the cell cycle phase at which genes reach peak transcription rate (green) and peak degradation rate (purple) for each pluripotent system. **c** Pairwise histograms of the circular correlation *r*_circ_ between transcription (green, upper triangle) and degradation (purple, lower triangle) timing for genes shared between each pair of pluripotent systems; red vertical lines indicate the median *r*_circ_ for each comparison, and n indicates the number of shared genes analyzed.

To resolve the cell cycle dependence of the underlying kinetic processes—transcription and degradation—we implemented a kinetic model based on RNA velocity [11, 38]. We used velocyto to obtain genome-wide quantification of unspliced and spliced mRNA, Sanity to obtain normalized, log-transformed values of each, and then solved an RNA velocity-based kinetic model to infer cell cycle-dependent transcription and degradation rates for each gene (see **Methods**). The bottom two rows of Fig. 3a show these cell cycle-dependent transcription (green) and degradation (purple) rates. Genes reached peak transcription and degradation in a specific, non-random order across the cell cycle, providing mechanistic insight into the kinetics of mRNA production and decay in each pluripotent system. Fig. 3b shows histograms of the timing of peak transcription (green) and peak degradation (purple) across genes; both processes showed prominent, organized waves across the cell cycle, again suggesting differences in transcriptional resource management across species and pluripotent states.

To perform a gene-by-gene comparison across systems, we calculated the circular correlation between the transcription and degradation rates across all combinations of the six pluripotent systems, restricting each comparison to genes shared between the corresponding pair of systems. Fig. 3c shows, for each pairwise comparison between systems, histograms of the circular correlation between transcription (green) and degradation (purple) timing; the red line marks the median circular correlation across genes for each comparison. Across nearly all pairwise comparisons, degradation timing was more highly correlated between systems than transcription timing, indicating that the kinetics of post-transcriptional regulation are more conserved across pluripotent systems than the kinetics of transcription itself.

Collectively, these kinetic analyses reveal that beneath the species- and state-specific differences in gene expression amplitude and phase distribution described above, the transcription and degradation programs driving the cell cycle are highly organized in time, unfolding as coordinated waves across the phase axis in every system examined. Critically, the timing of mRNA degradation is substantially more conserved across pluripotent systems than the timing of transcription, suggesting that post-transcriptional regulation—rather than transcriptional control—constitutes the more evolutionarily and developmentally stable layer of cell cycle regulation in pluripotent cells.

### Cell cycle regulators show increase in the amplitude of oscillations during progression from hiPSCs to endoderm

To study the changes in cell cycle regulation during development, we used a 3-day protocol (Fig. 4a; see **Methods**) to differentiate hiPSCs into definitive endoderm cells. Fig. 4b shows a principal component analysis using the top 100 highly variable genes across the three combined time points. The first PC captured transcriptional differences between day 1 and day 3, as well as between day 2 and day 3, while the second PC captured differences between day 1 and day 2. As cells progressed toward the endoderm lineage, expression of the pluripotency marker *POU5F1* (also known as *OCT3/4*) gradually declined, while expression of *SOX17*, a marker of definitive endoderm, progressively increased. Fig. S6 shows expression levels of additional pluripotency and definitive endoderm markers.

**Figure 4.**
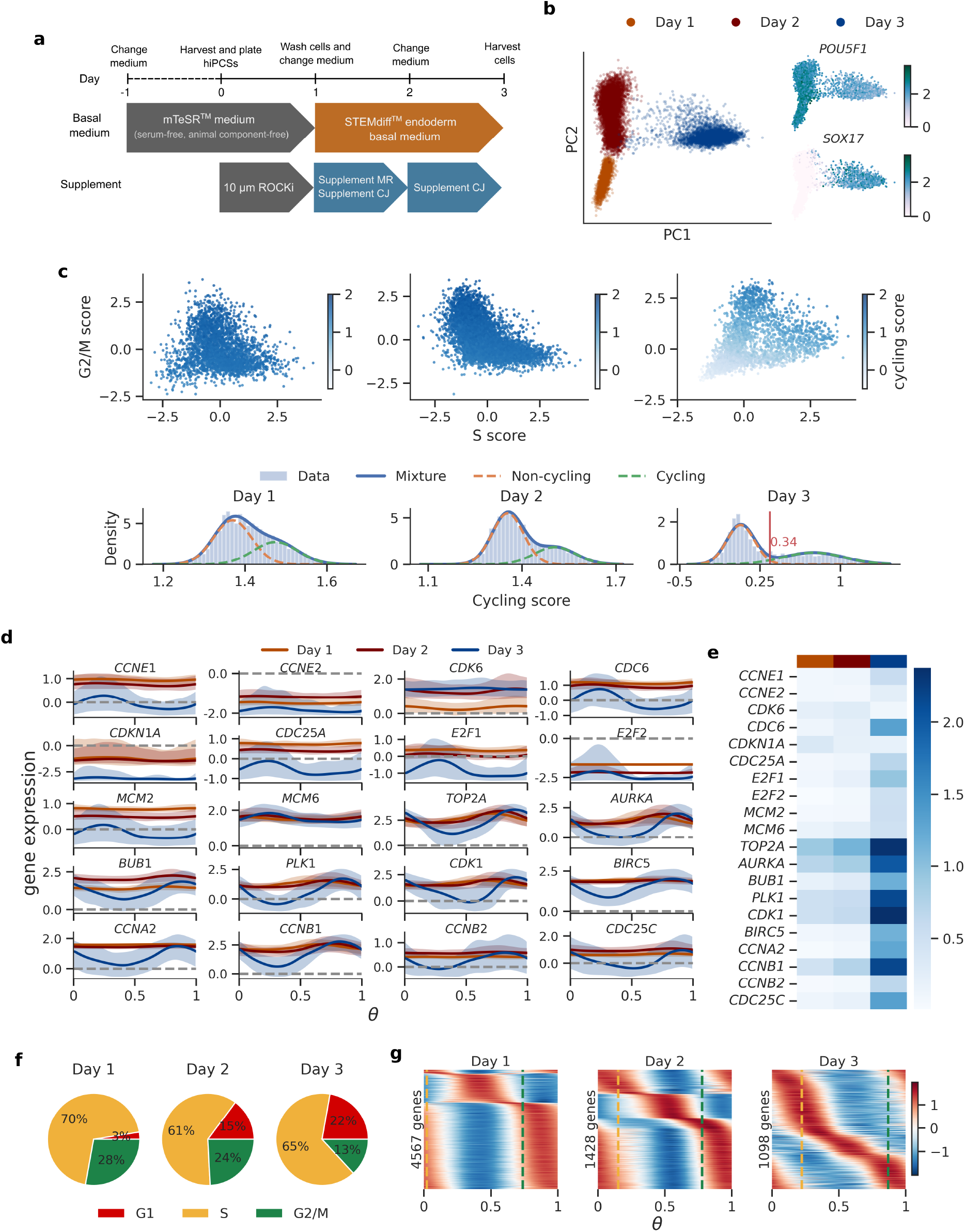
Cell cycle dynamics during commitment to the endoderm lineage. **a** Schematic of the 3-day directed differentiation protocol used to convert hiPSCs into definitive endoderm cells, indicating basal medium and supplement changes at each day (see Methods). **b** Left: cells from days 1 (orange), 2 (maroon), and 3 (blue) projected onto the top two principal components of gene expression. Right: expression of the pluripotency marker *POU5F1* and the definitive endoderm marker *SOX17* projected onto the same PC space. Additional pluripotency and endoderm marker genes are shown in Fig. S6. **c** Top: cells at each time point projected onto the S–G2/M manifold and colored by cycling score. Bottom: distribution of cycling score across all cells at each time point (light blue bars), with a fitted two-component Gaussian mixture model (blue) decomposed into non-cycling (orange dashed) and cycling (green dashed) components; the red line at day 3 indicates the threshold (0.34) used to exclude non-cycling cells from downstream phase inference. **d** Expression dynamics of 20 representative cell cycle regulators as a function of inferred cell cycle phase *θ* at day 1 (orange), day 2 (maroon), and day 3 (blue); shaded regions indicate confidence intervals. **e** Heatmap of oscillation amplitude for each cell cycle regulator (rows) at each time point (columns). **f** Percentage of cells in G1 (red), S (yellow), and G2/M (green) phase at each time point, based on phase boundaries shown in Fig. S7. **g** Heatmaps of phase-ordered spliced gene expression for the top oscillatory genes at each time point, filtered by explained variance and amplitude criteria (number of genes indicated above each panel); dashed yellow and green lines mark the inferred G1/S and S/G2 phase boundaries.

While performing cell cycle phase inference at each time point, we noted a substantial population of cells with low S and G2/M scores, suggesting these cells were either quiescent or spending an extended period in G1. To exclude quiescent cells from phase inference, we assigned a “cycling score” to every cell at each time point (see **Methods**). The top row of Fig. 4c visualizes cycling score on the S–G2/M manifold at days 1, 2, and 3; a substantial fraction of day 3 cells show low cycling scores. The bottom row of Fig. 4c shows the distribution of cycling scores across all cells at each time point. Fitting a two-component Gaussian mixture model to these distributions revealed pronounced bimodality at day 3, indicating that a significant fraction of cells likely exited the cell cycle and became quiescent or senescent. Because accurate phase inference requires that cells be actively cycling, we set a threshold at the intersection of the two Gaussian components (0.34) and retained only cells above this threshold for phase inference at day 3. Bimodality was already detectable at day 2, but was far less pronounced than at day 3, and overall cycling scores at days 1 and 2 were substantially higher than at day 3; we therefore included all cells at these two earlier time points in phase inference. Table S3 enlists genes that were highly expressed in the “non-cycling” endodermal cells, these could serve as markers of quiescence in human pluripotent systems.

Fig. 4d shows expression dynamics of cell cycle regulators at day 1 (orange), day 2 (maroon), and day 3 (blue). Cell cycle regulators were generally maintained at higher expression levels at days 1 and 2 than at day 3. Fig. 4e shows a heatmap of oscillation amplitude at each time point, revealing a marked increase in amplitude by day 3. Together, these results suggest that at earlier time points, cell cycle regulators oscillate around a comparatively high, sustained baseline, whereas by day 3 the larger oscillation amplitudes suggest a transition in cellular resource allocation—away from maintaining consistently high cell cycle regulator expression and toward the metabolic demands of establishing endodermal identity.

Fig. 4f shows the fraction of cells in G1 (red), S (yellow), and G2/M (green) phases, based on phase transition boundaries defined in Fig. S6. As cells progressed from hiPSCs to endoderm, the fraction of cells in G1 increased while the fractions in S and G2/M correspondingly decreased. Finally, Fig. 4g shows heatmaps of oscillatory expression dynamics for the top cycling genes at each time point. Using the same explained-variance and amplitude criteria as before, we found distinct, organized waves of gene expression across the cell cycle at all three time points, though the number of oscillatory genes declined sharply over the course of differentiation: 4,567 genes at day 1, 1,428 at day 2, and 1,098 at day 3. Fig. S7 shows the corresponding enrichment analysis for biological processes active during G1, S, and G2/M. Beyond the expected processes—DNA replication, ribosome biogenesis, protein translation, mitotic spindle organization, and nuclear division—we found enrichment for DNA repair during G1 and for protein phosphorylation, GTPase activity, and regulation of signal transduction during G2/M.

Together, these results show that the transition from pluripotency to definitive endoderm is accompanied by a coordinated remodeling of cell cycle behavior: a growing fraction of cells exit active cycling altogether, the surviving cycling population shifts toward a longer G1 phase, and the cell cycle regulators that remain oscillatory do so with markedly larger amplitude and across a shrinking set of genes. Rather than simply slowing down, differentiating cells appear to reallocate transcriptional and metabolic resources away from sustaining high baseline expression of cell cycle machinery and toward the acquisition of endodermal identity—suggesting that cell cycle remodeling is not a passive byproduct of differentiation but an actively regulated component of the lineage transition itself.

## Limitations of the study

Several limitations should be noted. First, pluripotent stem cell systems are cultured under artificial, defined conditions and thus represent approximations of the in vivo naїve and primed epiblast states rather than faithful recapitulations of them. Relatedly, growth medium alone can substantially shape transcriptional and cell cycle phenotypes: mESCs cultured in 2i+LIF versus LIF are nominally the same cell type, yet show distinct noise characteristics and cell cycle kinetics (Figs. 1b, 2f–h). Because culture condition, species, and developmental state are not fully decoupled across our panel, some of the differences we attribute to species or naїve/primed identity may be partly confounded by media effects.

Second, our kinetic model inherits known limitations of the RNA velocity framework. Unspliced counts are much lower than spliced counts, limiting statistical power for rate estimation, and standard oligo-dT-primed library preparation introduces 3^*′*^ capture bias that can distort unspliced-to-spliced ratios in a length- and isoform-dependent manner. These factors likely affect the inferred transcription and degradation rates across systems and could contribute to the overall lower correlation between transcription rates.

Finally, all cell cycle phase and kinetic estimates reported here are transcriptional, derived from mRNA abundance alone. They do not capture protein-level dynamics, shaped by translation, protein stability, and post-translational regulation, which are known to play a central role in cell cycle control. The oscillatory patterns described here therefore reflect the transcriptional layer of cell cycle regulation and may not directly correspond to the corresponding protein dynamics.

## Discussion

Pluripotent stem cell systems are a powerful tool for dissecting gene function and the physiological processes underlying development. Beyond providing scientific and mechanistic insight into cell fate decisions during early development—such as gastrulation—they also offer promising avenues for translational research, including regenerative medicine such as beta-cell replacement therapy. In this work, we compared transcriptional profiles, gene expression dynamics, and mRNA kinetics across the cell cycle in six pluripotent systems spanning two species (mouse and human) and both pre- and post-implantation epiblast states. Our analysis uncovered pronounced transcriptional divergence and distinct cell cycle dynamics across these systems, revealing fundamental differences in their underlying biology. We further characterized how cell cycle dynamics change as cells exit pluripotency and commit to a specific lineage, focusing on the transition from hiPSCs to definitive endoderm.

It is well known that scRNA-seq are extremely noisey due to intrinsic biological fluctuations and measurement noise. Sanity removes this confounding noise and renders gene expression variability—particularly for lowly expressed genes—independent of the measurement depth [39]. Sanity’s mean-centered expression values allow for direct cross-species comparison of marker genes: we found that canonical pluripotency markers such as *NANOG, POU5F1* (*OCT3/4*), and *MYC* were expressed at comparatively low levels in several systems, raising the possibility that their protein levels are buffered and remain relatively stable despite lower transcript abundance. We also observed inter-species differences, as well as differences between naїve and primed states, in the fraction of unspliced versus spliced mRNA—an observation consistent with divergence in splice variant and isoform usage during early embryonic development. And more generally, we found differences in transcriptional complexity across the pluripotent systems, most notably between mESCs, hESCs, and hiPSCs, which share broadly similar noise characteristics. Consistent with this, marker-space PCA revealed continuous, overlapping distributions of cells across systems rather than sharply isolated clusters (Fig. 1f), and individual markers such as *KLF2, PRDM14*, and *ZFP42* did not partition cleanly along expected naїve/primed lines (Fig. 1e), together suggesting that naїve and primed identity are better understood as a continuum of pluripotent states than as two discrete, categorically distinct programs.

Trajectory inference is used to order single-cell omics data along a path that reflects a continuous transition between cells. In this work we introduced the S-G2/M manifold, which allows to obtain a rough heuristic for the cell cycle dynamics as viewed from a transcriptomic perspective. We also developed Ciclopes, a deep learning tool that jointly infers the cell cycle phase with a neural network, the gene expression dynamics with a Fourier-based model, and the underlying mRNA kinetics, including the cell cycle dependence of transcription and degradation. Applying this framework, we found that the fraction of cells in G1, S, and G2/M varied systematically across pluripotent systems, with primed systems generally showing a more pronounced G1 phase. Cell cycle regulators showed higher mean expression in mouse systems than in human systems, but human systems showed larger oscillatory amplitude around their (lower) baseline. A similar pattern emerged during differentiation: as hiPSCs progressed toward definitive endoderm, cell cycle regulators showed reduced baseline expression but increased oscillatory amplitude. Together, these parallel trends—across species and across differentiation—suggest that as cells become more specialized, whether through evolutionary divergence or lineage commitment, they impose tighter, more sharply time-restricted control over cell cycle regulator expression, potentially to accommodate the metabolic demands of species- or lineage-specific gene programs. In contrast, correlation analysis between transcription and degradation timing revealed that post-transcriptional kinetics remain considerably more conserved across pluripotent systems than transcriptional kinetics themselves, pointing to mRNA degradation as a more evolutionarily stable layer of cell cycle control. Further experimental validation—particularly at the protein level—will be needed to test these hypotheses directly.

Taken together, our results establish that cell cycle regulation in pluripotent stem cells is not a fixed, universal program but a tunable one—shaped by species identity, developmental state, and lineage commitment, and implemented through distinct combinations of baseline expression and oscillatory amplitude rather than a single conserved strategy. The relative conservation of post-transcriptional kinetics amid this variability suggests a layered model of cell cycle control, in which mRNA degradation dynamics form a more stable substrate upon which more plastic, context-dependent transcriptional programs are overlaid. More broadly, this work underscores the importance of benchmarking pluripotent model systems against one another before treating them as interchangeable proxies for early mammalian development, and provides a quantitative frame-work—Ciclopes—for doing so in future studies of cell cycle-coupled differentiation and disease.

## Data and code availability

https://doi.org/10.5281/zenodo.21809002

https://github.com/mauliknariya/Ciclopes

## Acknowledgments

We are grateful to Gérard Gradwohl, Valérie Schreiber, and Stephane Vincent for guidance with hiPSC differentiation protocol, constructive criticism, and a thorough review of the manuscript. This work of the Interdisciplinary Thematic Institute IMCBio+, as part of the ITI 2021–2028 program of the University of Strasbourg, CNRS, and INSERM, was supported by IdEx Unistra (ANR-10-IDEX-0002) and by the SFRI-STRAT’US project (ANR-20-SFRI-0012) and EUR IMCBio (ANR-17-EURE-0023) under the framework of the France 2030 Program. Library preparation and sequencing were performed by the GenomEast platform, a member of the France Génomique consortium (ANR-10-INBS-0009).

## Author contributions

Conceptualization, N.M. and M.K.N.; methodology,

N.M. and M.K.N.; formal analysis, M.K.N., G.Z., and I.K.B.; investigation, D.S.-A. and C.T.-C.; data curation, M.K.N. and T.Y.; software, M.K.N.; visualization, M.K.N.; writing – original draft, M.K.N. and N.M.; writing – review & editing, D.S.-A., G.Z., S.J.; funding acquisition, N.M. and S.J.; project administration, N.M. and M.K.N.; supervision, N.M.

## Methods

### Culture of hiPSC

hiPSC line SB AD3.1 was maintained undifferentiated in mTeSR1 medium (STEMCELL™ Technologies). Cells were seeded in a Cultrex-coated p35 plate and split using mechanical picking and clump passaging every 3 or 4 days.

Differentiation to definitive endoderm Definitive endoderm was generated using STEMdiff™ Definitive Endoderm kit (STEMCELL™ Technologies). Cells were harvested with TrypLE select (Thermo Fisher) and seeded at 2 *×* 10^6^ on Cultrex-coated p35 plates containing mTeSR1 medium supplemented with 10*µ*M Y27632 (STEMCELL™ Technologies) for 24 hours (day 1). Plates were washed the next day with DMEM/F12 with 15mM HEPES and cultured in STEMdiff™ Endoderm Basal medium, supplemented with MR and CJ supplements (STEMCELL™ Technologies) for 24 hours (day 2). The following day, plates were washed with DMEM/F12 with 15mM HEPES and cultured in STEMdiff™ Endoderm Basal medium, supplemented with CJ supplement for 24 hours (day 3).

### Flow cytometry analyses

Cells were harvested with TrypLE select, quenched with DMEM/F-12 with 15 mM HEPES and washed twice with DPBS. Cells were fixed with ice-cold 70% ethanol for 1 hour at −20°C. Cells were pelleted at 500 *×*g, resuspended in wash buffer (PBS and 2% FBS) and incubated at room temperature for 5 minutes. Cells were washed two more times, then resuspended in PBS supplemented with 0.5% BSA and incubated with the fluorochrome-conjugated primary antibodies anti-Sox17 and anti-Oct3/4 (BD Pharmingen) for 1 hour at 4°C. Cells were washed twice and resuspended at 1*×*10^6^ cells/mL in PBS with 1% BSA. Finally, cells were filtered on 85 *µ*m nylon mesh and analyzed on BD Fortessa LSR II Cell analyzer.

### Single cell RNA-seq preparation and sequencing

At each time point, cells were harvested using TrypLE Select, quenched with DMEM/F12 supplemented with 15 mM HEPES, washed twice with DPBS, and resuspended in DPBS supplemented with 0.04% BSA. Library preparation was performed at the GenomEast platform of the Institut de Génétique et de Biologie Moléculaire et Cellulaire (IGBMC). Cells were loaded onto the 10X Genomics Chromium Controller with a target recovery of 7,000 single cells. 3^*′*^ gene expression (GEX) libraries were generated using the Chromium GEM-X Single Cell 3^*′*^ Kit according to the manufacturer’s instructions. GEX libraries were sequenced on an Illumina NextSeq 2000 using paired-end 28 + 85 bp reads. Image analysis and base calling were performed using Real-Time Analysis (RTA) software version 2.7.7 and BCL Convert version 3.8.4, respectively.

### Pluripotent systems

The details of cell types and culture conditions of the pluripotent systems which were not produced this work can be found in these sources:

| System/Reference | GEO accession |
| --- | --- |
| mESCs(2i+LIF)<br>Riba et al [10] | GSE167609 |
| mESCs(LIF)<br>Hu et al [15] | GSE264348 |
| mEpiSCs<br>Tomoda et al [16] | GSE166066 |
| hNES1<br>Taubenschmid-Stowers et al [17] | GSE178379 |
| hESCs<br>Han et al [18] | GSE157773 |
| hiPSCs<br>This work | GSEXXXXXX |

### Shannon’s diversity index

We used Shannon’s diversity index to quantify the transcriptional complexity in the gene expression of the model systems. For a given cell we define the Shannon’s diversity index as,

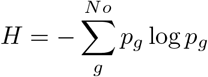

where *p*_*g*_ denotes the fraction of counts coming from gene *g*. To account for the differences in number of genes in the six data data sets, we calculated the Shannon index using the genes that were common between the six datasets, *N*_genes_ = 8, 166. Interpretation of the Shannon index: if all the counts in the cell are coming from one gene then the Shannon index would be, *H* = *−*(1) *×* log(1) = 0, whereas if the counts were uniformly distributed across 8,166 genes then the Shannon’s diversity index would be, 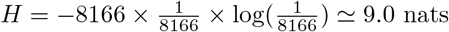. In general,

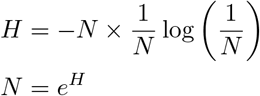

where *N* denotes the estimated number of genes contributing to signal (assuming that all the counts were uniformly distributed across all the genes).

### Data Processing

We used CellRanger v9.0.1 to process the fastq files and obtain the feature *×* barcode (or the gene *×* cell) matrix. We then used velocyto v0.17.17 to map the reads to the intronic and the exonic regions and obtain the unspliced and spliced counts. We used Sanity, a probabilistic modeling framework based on Bayesian statistics, to denoise the counts of gene counts as well as the unspliced and spliced (three independent runs of Sanity for each matrix). We use the log-transformed quotients (LTQs) of gene counts, unspliced, and spliced counts for all downstream computational analysis and modeling. When comparing between pluripotent systems we mean-centered the LTQs by subtracting the mean LTQs across all the cells and all the genes in a given dataset.

### Ciclopes

Ciclopes is a biology-informed deep learning tool for inferring the transcriptional cell cycle phase and mRNA kinetics from single-cell resolution data. It has two main components, the Phase model and the Kinetic model

#### Phase model

The Phase model is based on the autoencoder (AE) architecture, where the encoder is a neural network and infers the continuous cell cycle phase, whereas the decoder is a Fourier model which fits an oscillatory expression dynamics. We use the S-G2/M manifold (as described in Fig. 2b) to obtain an initial estimate of the cell cycle phase. In particular, we map every point on this manifold onto the red loop, which is the line of median density and represents the cell cycle dynamics on this manifold, and obtain an initial estimate of the cell cycle phase, *θ*_init_. Using this initial estimate we filter genes with an oscillatory dynamics based on their explained variance and amplitude over baseline gene expression (see the subsection on **Filtering genes with oscillatory profiles**). We then fit the Fourier model to these genes and obtain the Fourier coefficients,

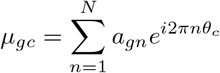

where, *µ*_*gc*_ : the predicted expression of gene *g* in cell *c, a*_*gn*_ is the *n*^th^ Fourier coefficients of gene *g, θ*_*c*_ is the cell cycle phase of cell *c*. We initialize the AE with the coefficients, *a*_*gn*_ and iteratively refine inference the cell cycle phase and the Fourier coefficients.

#### Kinetic model

The kinetic model is a system of coupled ordinary differential equations in unspliced, *u*_*g*_ and spliced, *s*_*g*_, such that transcription rate, *α*_*g*_, and degradation rate, *γ*_*g*_, are cell-cycle dependent, whereas the splicing rate, *β*_*g*_, is cell-cycle independent.

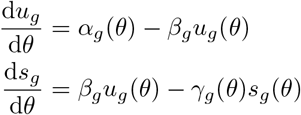

This can be solved pseudoanalytically using Fourier series (see supplemental information of Nariya et al. [11]). Note that owing to the sparsity in the unspliced quantifications, we *z*-scored the *u* and *s* values for every gene in the current implementation.

#### Loss terms

Since Sanity removes the Poissonian noise in the data, we assume that the predicted gene expression states are normally distributed around the observed gene expression values, i.e. the reconstruction loss is given by,

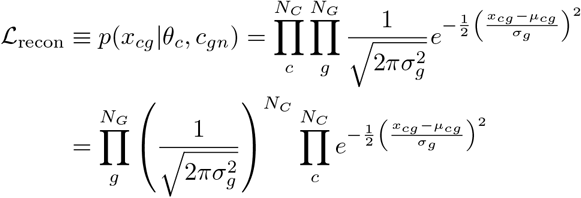

Integrating over all possible values of *σ*_*g*_, assuming assume a flat prior on *σ*_*g*_

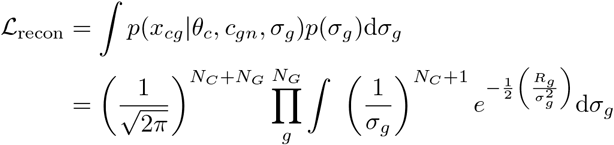

where 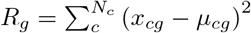

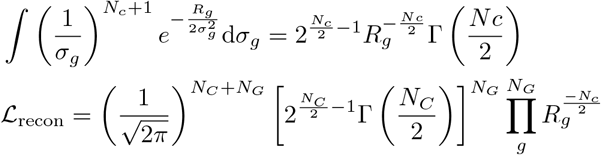

Finally, the reconstruction loss can be written as

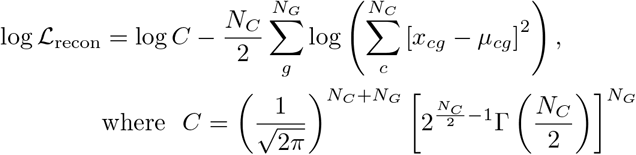

To ensure the inferred phases *θ* spread uniformly around the circle (avoiding mode collapse), we add a circular entropy regularization term. We construct a soft circular histogram of the batch’s *θ* estimates by kernel-smoothing with a von Mises kernel, the circular analog of a Gaussian, centered at each histogram bin, and maximize the (normalized) Shannon entropy of the resulting distribution.

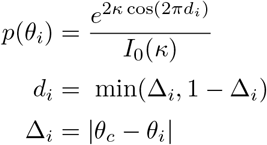

where *I*_0_ is the modified Bessel function of the first kind of order 0, Δ_*c*_ is the difference between the inferred theta, *θ*_*c*_ and the bin centers *θ*_*i*_, and *d* is the circularized distance.

The circular entropy is,

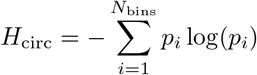

Lastly, to keep the inferred *θ*s close the initial estimates which are obtained from known biology, we introduce an “anchor” loss term,

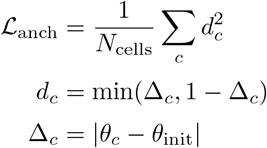

Altogether, total loss is given by,

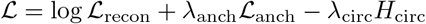

### Filtering genes with oscillatory profiles

We used the explained variance criterion and amplitude of oscillations to determine the genes with oscillatory expression dynamics. We examined the gene expression with respect to *θ*, and obtained their moving averages, *x*^ker^, using a periodic Gaussian kernel. For every gene we calculated the explained variance, *r*^2^

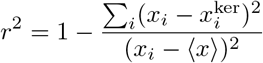

We also calculated the fold change in amplitude over the baseline gene expression for every gene,

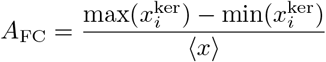

We filtered genes that satisfied this condition

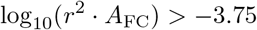

### Determination of cell cycle phase boundaries

In order to determine the M-G1 phase boundary, we used the mRNA counts in cells and fitted a monotonically increasing function (using a radial basis with an exponential kernel) with respect to *theta*, and shifted the *θ*s such that *θ* = 0 corresponds to the beginning of G1 phase. We used markers of G1–S and S-G2/M and assigned gene scores to cells for these transitions, we performed kernel smoothing of these scores with respect to *θ* and used the *θ* with maximum score to determine the G1–S and S–G2/M phase boundaries (details in https://github.com/mauliknariya/Ciclopes).

## Supplemental information

**Table S1:** Pluripotency markers. Table adapted from A Guide to Pluripotency Markers.

| Gene | Marker type | Molecular type | Species | Reference |
| --- | --- | --- | --- | --- |
| *ADCYAP1R1* | hPSC | Cell surface protein | Human | [26] |
| *ADGRG2* | Naïve, Primed | Cell surface protein | Human | [23, 24, 27] |
| *ATP1A1* | hESC | Cell surface protein | Human | [26] |
| *ATP1B3* | hESC | Cell surface protein | Human | [26] |
| *B3GAT1/CD57* | Primed | Cell surface protein | Human | [20, 23–25] |
| *BSG/CD147* | hESC, hPSC | Cell surface protein | Human | [26] |
| *CD7* | Naïve | Cell surface protein | Human | [20, 23–25, 27] |
| *CD24* | Naïve, Primed | Cell surface protein | Human | [20, 23–27] |
| *CD24* | Naïve, Primed | Cell surface protein | Human | [20, 23–25, 27] |
| *CD77* | Naïve, hESC | Cell surface antigen | Human | [20, 23–27] |
| *CD320* | Naïve | Cell surface protein | Human | [20, 27] |
| *CKAP4* | hESC | Cell surface protein | Human | [26] |
| *CRIPTO/TDGF1* | Regulation | Cell surface protein | Human | [25, 28] |
| *DDOST* | hESC | Cell surface protein | Human | [26] |
| *DNMT3L* | Naïve | Transcription factor | Human | [20, 25] |
| *DPPA2* | Naïve, Regulation | Transcription factor | Mouse | [21, 25] |
| *DPPA3* | Naïve | Transcription factor | Human | [20, 21, 23, 25] |
| *DPPA5* | Naïve | Transcription factor | Human | [20, 21, 25] |
| *DUSP6* | Primed | Transcription factor | Human | [20, 23] |
| *FNA3* | hPSC | Cell surface protein | Human | [26] |
| *EOMES/TBR2* | Regulation | Transcription factor | Mouse, Human | [21, 25] |
| *EPCAM/CD326* | hESC | Cell surface protein | Human | [26] |
| *ERBB2* | hESC | Cell surface protein | Human | [26] |
| *ERBB4* | hESC | Cell surface protein | Human | [26] |
| *F2* | hPSC | Cell surface protein | Human | [26] |
| *F11R* | Naïve, Primed | Cell surface protein | Human | [23, 24, 27] |
| *FAM216A* | hPSC | Cell surface protein | Human | [26] |
| *FGFR3* | hPSC | Cell surface protein | Human | [26] |
| *GATA6* | Naïve, Primed | Transcription factor | Mouse, Human | [21, 25] |
| *PODXL/GCTM2* | hESC, iPSC | Cell surface protein | Human | [27, 28] |
| *GLG1* | hESC | Cell surface protein | Human | [26] |
| *GPC4* | hESC | Cell surface protein | Human | [26] |
| *HLA-A* | Primed | Cell surface protein | Human | [20, 24] |
| *HLA-B* | Primed | Cell surface protein | Human | [20, 24] |
| *HLA-C* | Primed | Cell surface protein | Human | [20, 24] |
| *HSPA8, HSC70* | hESC | Cell surface protein | Human | [26] |
| *HTR2C* | hPSC | Cell surface protein | Human | [26] |
| *IL17RD* | hPSC | Cell surface protein | Human | [26] |
| *IL6ST, CD130* | Naïve | Cell surface protein | Human | [20, 23–25, 27] |
| *ITGAV* | hESC | Cell surface protein | Human | [26] |
| *KLF17* | Naïve | Transcription factor | Human | [20, 23, 25] |
| *KLF4* | Reprogramming | Transcription factor | Mouse, Human | [20, 21, 23–25, 27] |
| *KLF5* | Naïve | Transcription factor | Human | [21, 23, 25] |
| *L1CAM/CD171* | hESC | Cell surface protein | Human | [26] |
| *LAMP2/CD107b* | Naïve | Cell surface protein | Human | [20, 24] |
| *LY9/CD229* | Naïve | Cell surface protein | Human | [20, 24] |
| *MYC* | Reprogramming | Transcription factor | Human | [24, 25, 27] |
| *NANOG* | Reprogramming | Transcription factor | Mouse, Human | [20, 21, 23–25, 28] |
| *NLGN4X* | Primed | Cell surface protein | Human | [23, 24, 26, 27] |
| *NPR1* | hPSC | Cell surface protein | Human | [26] |
| *OPCML* | hPSC | Cell surface protein | Human | [26] |
| *OTX2* | Primed | Transcription factor | Mouse, Human | [20, 21, 23] |
| *PCDH1* | Naïve, Primed | Cell surface protein | Human | [23, 24, 27] |
| *PODXL* | hESC | Cell surface protein | Human | [26] |
| *POU5F1/OCT4* | Reprogramming | Transcription factor | Mouse, Human | [20, 21, 23–28] |
| *PTPRZ1/PTPRZ* | hESC | Cell surface protein | Human | [26] |
| *RTN3* | hESC | Cell surface protein | Human | [26] |
| *RTN4* | hESC | Cell surface protein | Human | [26] |
| *SIRPA/CD172a* | Primed, hESC | Cell surface protein | Human | [23, 24, 26] |
| *SLC7A5* | hESC | Cell surface protein | Human | [26] |
| *SOX2* | Reprogramming | Transcription factor | Mouse, Human | [20, 21, 23–28] |
| *SSEA-1* | Naïve, Primed | Cell surface antigen | Mouse | [21, 28] |
| *SSEA-3* | Naïve, Primed | Cell surface antigen | Human | [24, 27, 28] |
| *SSEA-4* | Naïve, Primed | Cell surface antigen | Human | [20, 23, 24, 27, 28] |
| *SSEA-5* | hESC | Cell surface protein | Human | [26, 28] |
| *ST6GAL1/CD75* | Naïve | Cell surface protein | Human | [20, 23–25, 27] |
| *STAT3* | Reprogramming | Transcription factor | Human | [23–25] |
| *SUSD2* | Naïve | Cell surface protein | Human | [27] |
| *TFCP2L1* | Naïve | Transcription factor | Human | [20, 21, 23] |
| *TFRC/TFR1/CD71* | hESC, hPSC | Cell surface protein | Human | [26] |
| *THY1/CD90* | Primed | Cell surface protein | Human | [20, 23–25, 27, 28] |
| *TRA-1-60* | Naïve, Primed | Cell surface antigen | Human | [20, 23, 24, 28] |
| *TRA-1-81* | Naïve, Primed | Cell surface antigen | Human | [20, 24, 28] |
| *VAPA* | hESC | Cell surface protein | Human | [26] |
| *ZDHHC13* | hESC | Cell surface protein | Human | [26] |
| *ZIC2* | Primed | Transcription factor | Human | [20, 23, 25] |

**Table S2:** Cell cycle durations in pluripotent systems.

| System | Total cycle | G1 | S | G2/M | Reference |
| --- | --- | --- | --- | --- | --- |
| mESCs (LIF) | 12–14 h | ~2 h | 8–9 h | ~2 h | Savatier 1994 [30]; Stead 2002 [31]; ter Huurne 2017 (BrdU/PI: 18% G1 / 72% S / 10% G2) [2]; Waisman 2019 [32] |
| mESC (2i+LIF) | 13.25–13.5 h | 2–2.5 h | 7–9 h | 1.5–2.5 h | Waisman 2019 [32], ter Huurne (42% G1 / 48% S / 7% G2) [2] |
| mEpiSCs | 16–20 h (inferred) | 5–7 h | 7–9 h | 3–4 h | No direct measurement published; Coronado 2013 establishes the direction (G1 lengthens ground → naive → primed) but not absolute values [3] |
| Naïve hESCs | 16–20 h (inferred) | 4–5 h | 9–10 h | 3–4 h | Ware 2014 [33], Ware 2023 [34] |
| hESCs | 15–16 h | 2.5–3 h | ~8 h | ~5h | Becker 2006 [35] |
| hiPSCs | 16–18 h | ~2.5 h | ~8 h | 5–6 h | Ghule 2011 [36], Hannah 2010 [37] |

**Table S3:** Top 51 genes that are differential expressed in non-cycling endodermal cells (day 3), determined using Wilcoxon rank-sum test, when compared to cycling cells at the same time point.

| Gene | z-score | Gene | z-score | Gene | z-score |
| --- | --- | --- | --- | --- | --- |
| *GATA6-AS1* | 23.50 | *AC010624.5* | 18.79 | *HCG14* | 18.64 |
| *MIR302CHG* | 22.54 | *C16orf54* | 18.78 | *AC106801.1* | 18.64 |
| *CCL2* | 20.50 | *LINC00479* | 18.78 | *FOXA1* | 18.64 |
| *LINC00467* | 19.60 | *AC129492.1* | 18.78 | *AC083841.1* | 18.64 |
| *PHKA2-AS1* | 19.33 | *AL022341.1* | 18.77 | *PCDHB11* | 18.63 |
| *EGFLAM* | 19.28 | *SRARP* | 18.75 | *AL133299.1* | 18.62 |
| *MYOF* | 19.19 | *AC044810.2* | 18.74 | *GALR3* | 18.62 |
| *AL035252.5* | 19.14 | *CALHM5* | 18.74 | *AL592078.2* | 18.62 |
| *HIC1* | 19.04 | *KIF25-AS1* | 18.72 | *PCDHB12* | 18.61 |
| *FSBP* | 18.98 | *PRND* | 18.71 | *ZNF467* | 18.61 |
| *AC084809.1* | 18.92 | *F7* | 18.71 | *Z93943.1* | 18.61 |
| *AC096746.1* | 18.85 | *RAX* | 18.69 | *AL022157.1* | 18.61 |
| *Z98745.1* | 18.83 | *RPS6KB2-AS1* | 18.69 | *AC004923.4* | 18.61 |
| *PARD3-AS1* | 18.83 | *PRSS57* | 18.68 | *AC010336.3* | 18.59 |
| *NYX* | 18.81 | *AC005632.6* | 18.65 | *KRT19* | 18.58 |
| *MZB1* | 18.80 | *CRYBA2* | 18.65 | *AC004156.1* | 18.58 |
| *AC091043.1* | 18.80 | *LINC01270* | 18.65 | *SERPINF2* | 18.57 |

**Figure S1:**
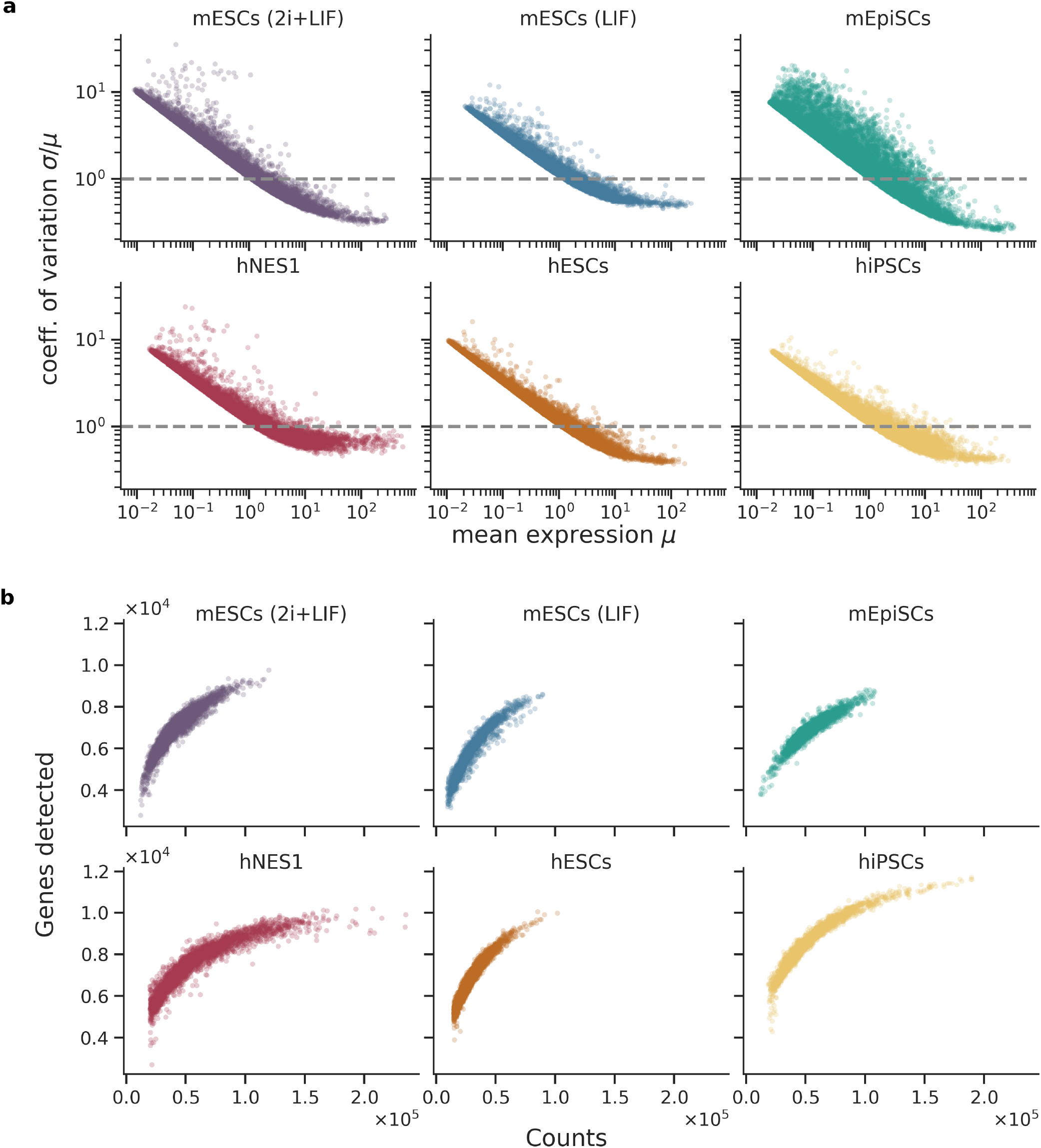
Noise characteristics and genes detected in the scRNA-seq data across model systems.

**Figure S2:**
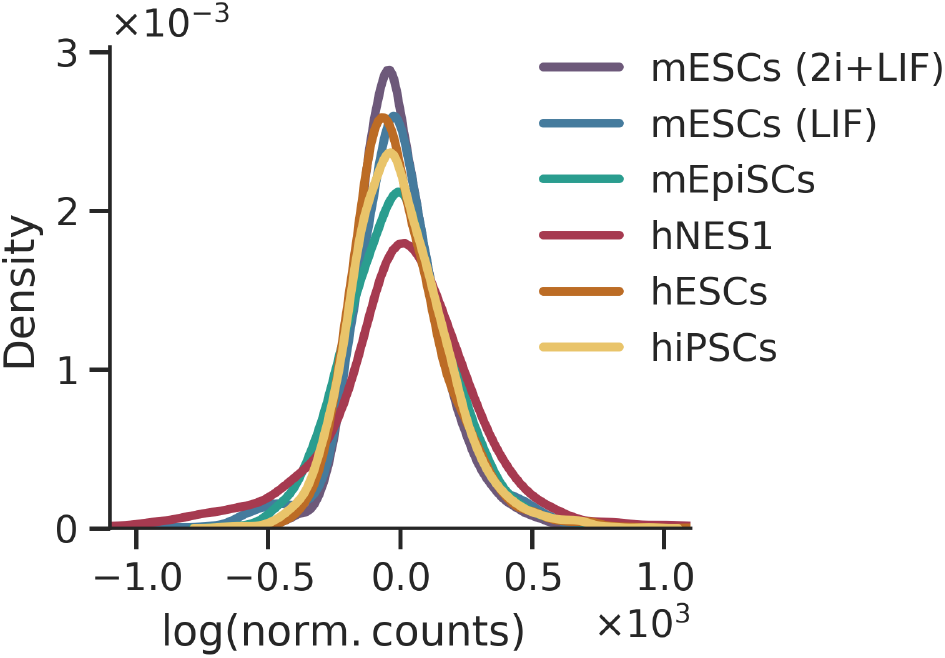
Distribution of mean-centered log-normalized counts in cells across the model systems.

**Figure S3:**
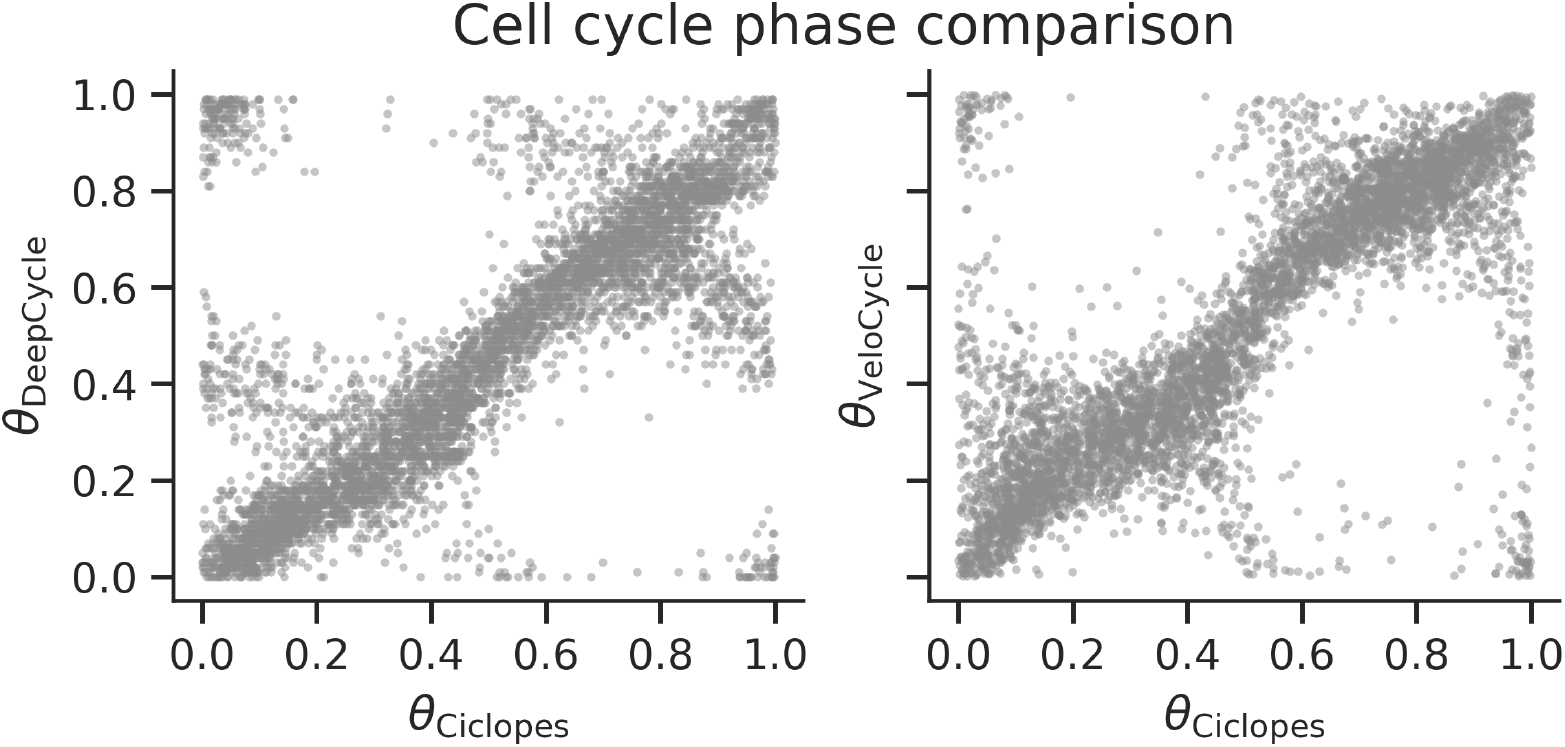
Comparison of cell cycle phases in mESCs (2i+LIF) inferred by Ciclopes, DeepCycle, and VeloCycle

**Figure S4:**
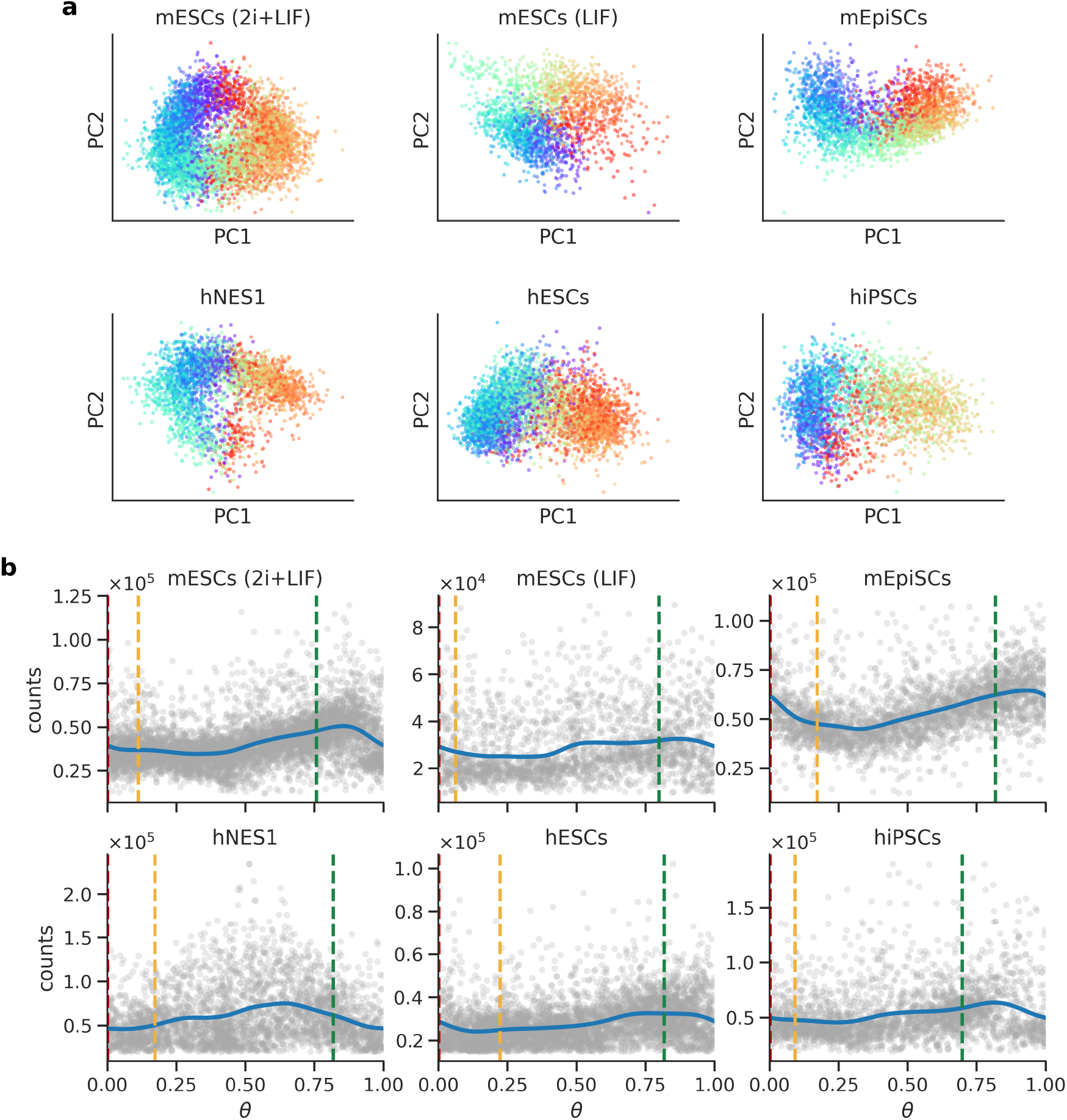
a: PCA showing the cell cycle dynamics across model systems. **b:** total mRNA counts versus the cell cycle phase *θ* for the different pluripotent systems.

**Figure S5:**
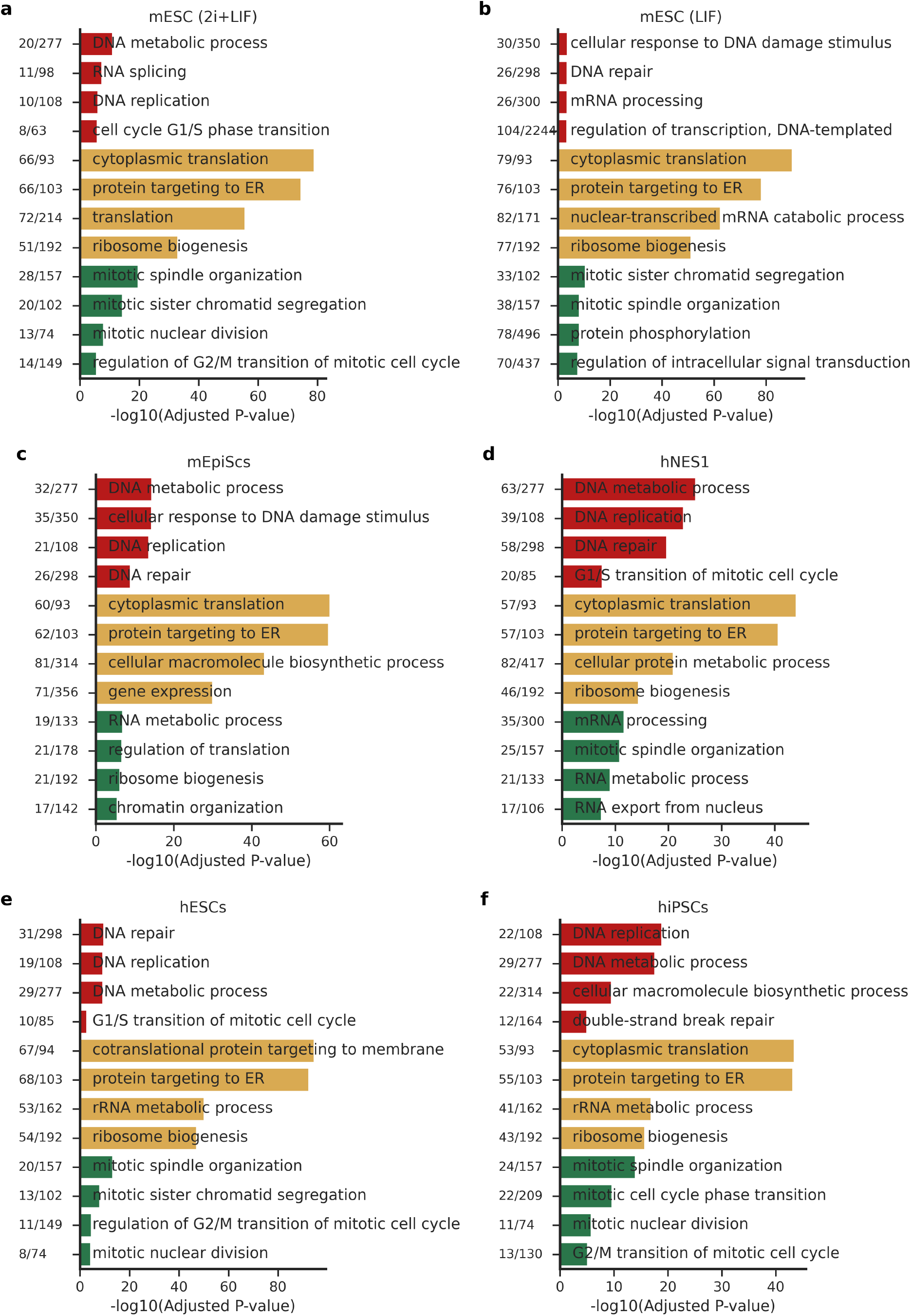
Enrichment analysis for genes with peakexpreesion in G1 (in red), S (in yellow) and G2/M (in green) phases.

**Figure S6:**
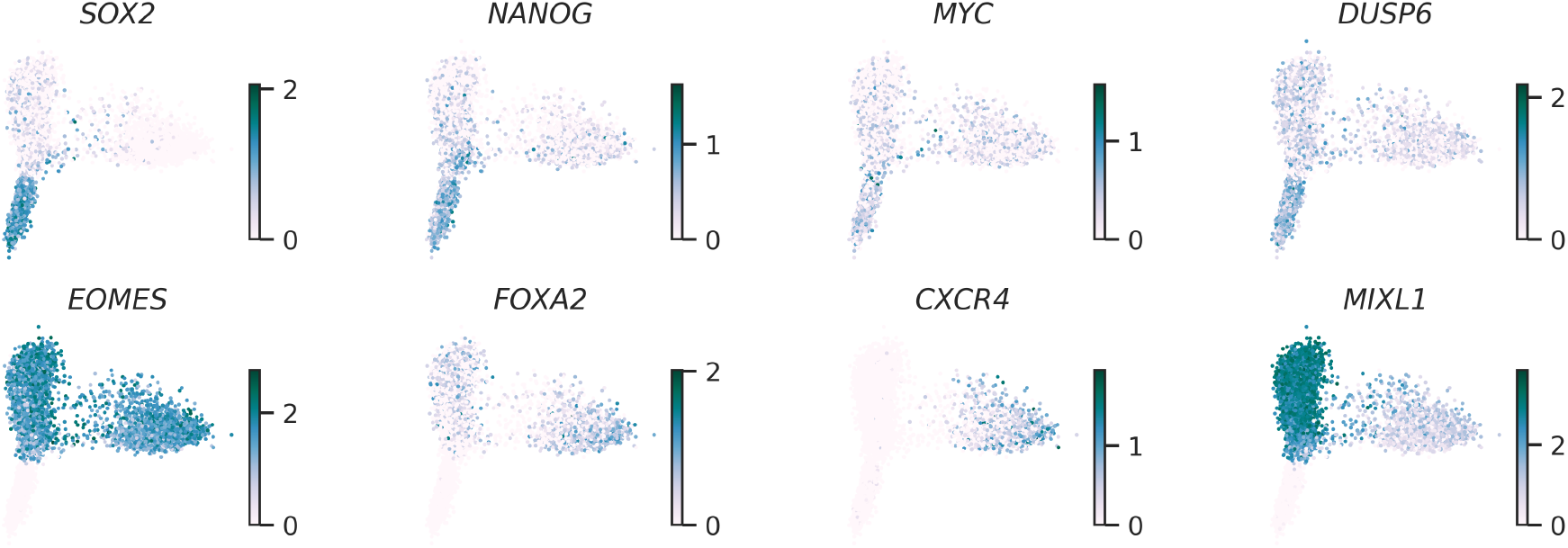
Gene expression of select plurioptency markers—*SOX2, NANOG, MYC*, and *DUSP6*, and that of the markers of definitive endoderm—*EOMES, FOXA2, CXCR4*, and *MIXL1*, visualized on the first two prinicpal components of the scRNA-seq profiles at the three timepoints.

**Figure S7:**
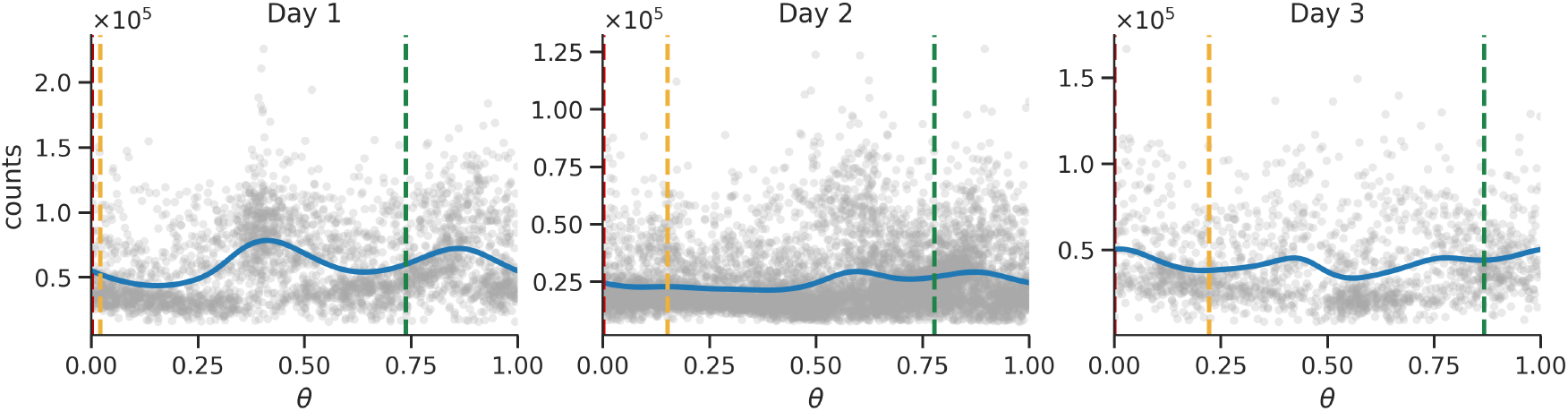
Total mRNA counts versus the cell cycle phase *θ* at day 1, 2, and 3.

## Notes

### Competing Interest Statement

The authors have declared no competing interest.

## References

1 B. Boward et al., “Concise review: control of cell fate through cell cycle and pluripotency networks”, Stem Cells 34 (2016).

2 M. ter Huurne et al., “Distinct cell-cycle control in two different states of mouse pluripotency”, Cell Stem Cell 21 (2017).

3 D. Coronado et al., “A short g1 phase is an intrinsic determinant of naïve embryonic stem cell pluripotency”, Stem Cell Research 10 (2013).

4 B. DeVeale et al., “G1/s restriction point coordinates phasic gene expression and cell differentiation”, Nature Communications 13 (2022).

5 S. Pauklin and L. Vallier, “The cell-cycle state of stem cells determines cell fate propensity”, Cell 155 (2013).

6 I. G. M. Brons et al., “Derivation of pluripotent epiblast stem cells from mammalian embryos”, Nature 448 (2007).

7 P. J. Tesar et al., “New cell lines from mouse epiblast share defining features with human embryonic stem cells”, Nature 448 (2007).

8 J. Nichols and A. Smith, “Naive and primed pluripotent states”, Cell Stem Cell 4 (2009).

9 K. C. Davidson et al., “The pluripotent state in mouse and human”, Development 142 (2015).

10 A. Riba et al., “Cell cycle gene regulation dynamics revealed by rna velocity and deep-learning”, Nature Communications 13 (2022).

11 M. K. Nariya et al., “Single-cell multiomics reveals the oscillatory dynamics of mrna metabolism and chromatin accessibility during the cell cycle”, Cell Reports 44 (2025).

12 S. Liang et al., “Latent periodic process inference from single-cell rna-seq data”, Nature Communications 11 (2020).

13 A. R. Lederer et al., “Statistical inference with a manifold-constrained rna velocity model uncovers cell cycle speed modulations”, (2024).

14 G. Zanardelli et al., “Spatiotemporal regulation of cell cycle states within the complex tumor microenvironment”, (2025).

15 X. Hu et al., “Artseq-fish reveals position-dependent differences in gene expression of micropatterned mescs”, Nature Communications 15 (2024).

16 K. Tomoda et al., “Reprogramming epiblast stem cells into pre-implantation blastocyst cell-like cells”, Stem Cell Reports 16 (2021).

17 J. Taubenschmid-Stowers et al., “8c-like cells capture the human zygotic genome activation program in vitro”, Cell Stem Cell 29 (2022).

18 M.-K. Han, “Single cell rna sequencing analysis of human embryonic stem cells, human embryoid bodies, human mesenchymal stem cells and human musculoskeletal stem cells”, Gene Expression Omnibus (2020).

19 Y. Chen and D. Lai, “Pluripotent states of human embryonic stem cells”, Cellular Reprogramming 17 (2015).

20 A. J. Collier et al., “Comprehensive cell surface protein profiling identifies specific markers of human naive and primed pluripotent states”, Cell Stem Cell 20 (2017).

21 S. Morgani et al., “The many faces of pluripotency: in vitro adaptations of a continuum of in vivo states”, BMC Developmental Biology 17 (2017).

22 R. M. Baldarelli et al., “Mouse genome informatics: an integrated knowledgebase system for the laboratory mouse”, Genetics 227 (2024).

23 A. J. Collier and P. J. Rugg-Gunn, “Identifying human naïve pluripotent stem cells-evaluating state-specific reporter lines and cell-surface markers”, BioEssays 40 (2018).

24 O. Trusler et al., “Cell surface markers for the identification and study of human naive pluripotent stem cells”, Stem Cell Research 26 (2018).

25 A. Yilmaz and N. Benvenisty, “Defining human pluripotency”, Cell Stem Cell 25 (2019).

26 A. Meyfour et al., “The quest of cell surface markers for stem cell therapy”, Cellular and Molecular Life Sciences 78 (2020).

27 J. Goodwin et al., “The application of cell surface markers to demarcate distinct human pluripotent states”, Experimental Cell Research 387 (2020).

28 P. W. Andrews and P. J. Gokhale, “A short history of pluripotent stem cells markers”, Stem Cell Reports 19 (2024).

29 W. Bouchereau et al., “Major transcriptomic, epigenetic and metabolic changes underlie the pluripotency continuum in rabbit preimplantation embryos”, Development 149 (2022).

30 P. Savatier et al., “Contrasting patterns of retinoblastoma protein expression in mouse embryonic stem cells and embryonic fibroblasts”, Oncogene 9 (1994).

31 E. Stead et al., “Pluripotent cell division cycles are driven by ectopic cdk2, cyclin a/e and e2f activities”, Oncogene 21 (2002).

32 A. Waisman et al., “Cell cycle dynamics of mouse embryonic stem cells in the ground state and during transition to formative pluripotency”, Scientific Reports 9 (2019).

33 C. B. Ware et al., “Derivation of naïve human embryonic stem cells”, Proceedings of the National Academy of Sciences 111 (2014).

34 C. B. Ware et al., “Derivation of naïve human embryonic stem cells using a chk1 inhibitor”, Stem Cell Reviews and Reports 19 (2023).

35 K. A. Becker et al., “Self-renewal of human embryonic stem cells is supported by a shortened g1 cell cycle phase”, Journal of Cellular Physiology 209 (2006).

36 P. N. Ghule et al., “Reprogramming the pluripotent cell cycle: restoration of an abbreviated g1 phase in human induced pluripotent stem (ips) cells”, Journal of Cellular Physiology 226 (2011).

37 J. H. Hanna et al., “Pluripotency and cellular reprogramming: facts, hypotheses, unresolved issues”, Cell 143 (2010).

38 G. La Manno et al., “Rna velocity of single cells”, Nature 560 (2018).

39 J. Breda et al., “Bayesian inference of gene expression states from single-cell rna-seq data”, Nature Biotechnology 39 (2021).

